# EEG Microstate Sequences as Potential Brain-Computer Interface Triggers Derived from Motor Imagery Classification

**DOI:** 10.64898/2026.08.18.745436

**Authors:** Annika Wollmann, Markus Goldhacker

## Abstract

EEG microstates are a distinct number of quasi-stable spatial distributions of brain activity. Microstate trajectories are strongly suspected to reflect the underlying neural mechanisms during information processing and are therefore also called the ”building blocks” of human thought. In this study, we examined, if EEG microstate sequences can serve as potential triggers for a Brain-Computer Interface (BCI). To this end, a semi-supervised deep learning model architecture consisting of an LSTM-based autoencoder and a dense neural network was utilized to classify between left- and right-hand motor imagery EEG data, with the resulting classification output serving as the BCI trigger. On the one hand, this was done in a 2-step approach, in which the autoencoder and classifer have been trained separately. On the other hand, an end-to- end approach was employed, where training was performed by combining reconstruction and classification losses. Results show that the proposed model architecture was able to extract relevant features from microstate sequences and exploit them for within subjects and sessions classification. Applying transfer learning to session-to-session or across-subject transfer resulted in peak classification accuracies around 89%. We also investigated to what extent transfer learning has to be applied to reach considerable classification accuracies serving as the calibration time representative. We found that on average around 400 s are needed for BCI calibration when emplyoing our approach to reach 80% classification accuracy. The present study signifies that the investigation of EEG microstate trajectories can be a promising approach for extracting BCI triggers, as it reduces the dimensionality of multi-channel recorded EEG signals to a distinct number of brain states over time. Deep learning methods, especially transfer learning, applied to EEG microstate trajectories seem promising regarding user-convenient and calibration-free BCIs in real-world applications.

## 1 INTRODUCTION

Since their introduction by Vidal (1973), BCIs, which enable human-computer interaction by utilizing measurable electrical brain signals, have become a well-established field of research across neuroscience, computer technology and engineering due to their promising applications. Controlling neuroprosthetic robotic limbs or spelling devices that transform brain signals into computerized speech are representative examples (Peksa and Mamchur, 2023). Challenges for creating a proper BCI include building up sufficient neurophysiological knowledge, applying appropriate signal analysis techniques to extract relevant features from brain signals, and perform classification of different mental tasks with sufficient accuracies. After the so-called offline training or calibration phase, a live application or online test phase is conducted that requires a functional and above all efficient computer science implementation. (Vidal, 1973; Maiseli et al., 2023; Peksa and Mamchur, 2023)

Brain signals for BCIs are predominantly measured using electroencephalography (EEG), due to its non-invasiveness, relative simplicity, and high temporal resolution (Peksa and Mamchur, 2023). Among EEG-based BCI paradigms, steady-state visual evoked potentials, P300, and Motor Imagery (MI) are most commonly used (Peksa and Mamchur, 2023; Aggarwal and Chugh, 2022), with the latter being the focus of this work. MI refers to the mental simulation of a movement without actual execution (Decety, 1996). It induces characteristic changes in brain activity that can be exploited to discriminate, e. g., between left- and right-hand movements (Pfurtscheller and Neuper, 2001).

However, the performance and generalizability of BCIs are still limited by challenges in EEG signal processing, including low signal-to-noise ratio, non-stationarity, and high inter-subject variability (Aggarwal and Chugh, 2022; Lotte et al., 2018; Roy et al., 2019). In particular, the high degree of heterogeneity between subjects explains why most BCIs are currently subject-specific, where training and application are performed on the same subject (Sartipi and Cetin, 2024; dos Santos et al., 2023). Reaching a sufficient amount of training data for one subject can be time-consuming and inconvenient, but generally yields high classification accuracy (Aggarwal and Chugh, 2022). In contrary, a subject-independent BCI, where subjects in training and testing phase differ, is desirable to achieve generalization across users but often leads to worse performances (Sartipi and Cetin, 2024; dos Santos et al., 2023). Despite considerable progress in recent years, no standard methodology has yet been established for developing robust subject-independent BCIs, limiting their applicability in real-world scenarios. In this context, recent work increasingly explores Machine Learning (ML) and Deep Learning (DL) approaches to address challenges such as manual feature extraction and inter-subject variability. In particular, Roy et al. (2019) highlight the growing emphasis on inter-subject learning using DL, with initial results showing modest performance gains over conventional EEG analysis methods and promising generalization capabilities.

Another line of research in the context of EEG focuses on EEG microstates, which are defined as quasi-stable spatial distributions of brain activity (Lehmann et al., 1998). It has been shown that only a small number of microstates can explain the majority of the continuous topographies of EEG signals (usually *>* 70%) and that in most cases the same four microstates predominate, therefore being referred to as the four canonical forms (A-D) (Khanna et al., 2015). These distinct spatial patterns, first introduced by Koenig et al. (1999), show a left-right orientation, a right-left orientation, an anterior-posterior orientation as well as a fronto-central maximum (Michel and Koenig, 2018), and seem to be caused by distinct settings of active neural populations and therefore are likely to reflect divergent functional brain states (Lehmann et al., 1987, 1998). The interpretation of microstates suggests that mental processs can be represented by a sequence of this limited number of microstates (Lehmann et al., 1987). Therefore, microstates are also called the ”homogeneous basic building blocks of brain information processing” (Lehmann et al., 1987) or the ”atoms of thought” (Lehmann et al., 1998).

Since studies combining the above mentioned lines of research are sparse, we examined whether and to what extent EEG microstate sequences can serve as potential BCI triggers in the MI paradigm. We define the resulting output of a left- and right-hand MI classification, which can be used to initiate subsequent BCI commands or actions, as a BCI trigger. To this end, EEG signals from nine subjects performing left- and right-hand MI each on two different days were obtained from the BCI Competition IV dataset 2a (Brunner et al., 2008). This data was used to extract microstate sequences and to further analyze their trajectories over time. Classification performance for the subject-specific and subject-independent BCI context was examined by applying a semi-supervised DL architecture. Furthermore, the study investigated the extent to which transfer learning is suitable for adapting pre-trained models to EEG data from new subjects or new recording sessions in the context of EEG microstate trajectories. In particular, our investigations culminated in an estimate of the calibration time required for adapting BCIs to unseen users.

## 2 RELATED WORK

In the following section, we elaborate in detail on the above mentioned research directions. Conventional feature extraction and classification methods are reviewed in the context of BCIs expanding them to deep learning approaches followed by a focus on subject-independent BCI strategies. We then examine a EEG microstate approach using aggregated temporal features for BCI classification, and detail on a study employing microstates in a non-BCI context, which is the substrate for our methodology transfering it to the BCI context.

### 2.1 Classification Approaches on Motor Imagery EEG Data

#### 2.1.1 Conventional Feature Extraction and Classification Methods

Conventional feature extraction methods rely on algorithmic approaches applied to preprocessed EEG signals, while still requiring a substantial degree of manual intervention. With regard to the dataset 2a of the BCI Competition IV, Common Spatial Pattern (CSP), that finds spatial filters of a bandpass-filtered EEG signal, is among the most widely used methods for deriving discriminating features of different MI tasks (Aggarwal and Chugh, 2019). All the solutions submitted in the BCI Competition were based on CSP, with the winning solution employing a Filter Bank CSP (FBCSP) algorithm (Ang et al., 2008; Tangermann et al., 2012), that applies the CSP method to a set of (non-overlapping) frequency bands (Ang et al., 2008). While the number of DL methods has increased in recent years, there are still studies that deal with the further development of CSP to improve classification performance, such as in the study of Das et al. (2020). The classification of the before-handed extracted features is usually done by the linear classifiers Linear Discriminant Analysis (LDA) and Support Vector Machine (SVM) or small neural networks (Lotte et al., 2018).

#### 2.1.2 Deep Learning Approaches

DL approaches have emerged in the past years to overcome the demanding step of manual feature engineering by automatically selecting relevant features. As DL techniques can be applied on raw or minimally preprocessed EEG data, discriminant patterns in all possible domains can be found by the ML algorithm and a manual decision of either one of them gets obsolete. Furthermore, the feature extraction and training of a classification algorithm can be done simultaneously and are therefore promising to yield more robust results than hand-crafted feature engineering (Lotte et al., 2018; Roy et al., 2019). However, DL methods typically require large amounts of training data due to the high number of trainable parameters, which poses a challenge for EEG-based applications where data availability is limited. This constraint is particularly critical in practical BCI systems, where short calibration times are desired (Lotte et al., 2018). A variety of DL architectures have been explored for EEG analysis, with Convolutional Neural Networks (CNNs), Recurrent Neural Networks (RNNs) and Autoencoders (AEs) being the most commonly used. CNNs are widely adopted due to their ability to capture hierarchical patterns and process raw EEG signals efficiently in an end-to-end manner. RNNs are particularly suitable for modeling temporal dependencies in non-stationary brain signals, whereas AEs are primarily used for dimensionality reduction and feature extraction, often employed in a two-step pipeline (Roy et al., 2019). To mitigate data limitations, shallower networks, regularization techniques, and data augmentation strategies such as overlapping window segmentation are often employed (Lotte et al., 2018; Roy et al., 2019; Schirrmeister et al., 2017). Overall, DL methods show slight performance improvement over traditional approaches and offer high flexibility for different tasks and datasets (Roy et al., 2019; Schirrmeister et al., 2017).

#### 2.1.3 Subject-Independent Approaches

Model performance of subject-independent BCIs is commonly evaluated using a Leave-One-Subject-Out (LOSO) cross-validation scheme, in which data from all except for one subjects is used for training and the remaining subject for testing. This procedure is repeated until each subject served once as the test set.

Lotte et al. (2009) investigated subject-independent BCI approaches on the dataset 2a of the BCI Competition IV. While FBCSP achieved the best performance for the subject-specific approach, it performed worst in the subject-independent case, likely due to the high heterogeneity between subjects in spatial and spectral distribution of discriminant segments of MI EEG signals. To address this, the authors proposed a multi-resolution decomposition combining broader and finer frequency bands to capture both subject-independent and subject-specific features, achieving an average accuracy around 71% for classifying left- and right-hand MI. (Lotte et al., 2009)

Luo et al. (2023) addressed subject-independent BCI classification by considering temporal variability of event-related desynchronization (ERD)/event-related synchronization (ERS) patterns in EEG signals. They proposed a CNN-based architecture with an attention mechanism to identify informative temporal segments in long EEG trials, followed by a small neural network classification block. Additionally, a mirror augmentation strategy exploiting contralateral ERD/ERS effects was introduced to increase the training data. Evaluated on dataset 2a, the method achieved an average accuracy of 67%, whereby the EEG signal was bandpass filtered to 8-30 Hz, reduced to the channels C3, Cz and C4 and a time window of 3.5 seconds (0.5-4 seconds after cue) was selected. (Luo et al., 2023)

Sartipi and Cetin (2024) proposed a semi-supervised model architecture for subject-independent classification, combining a CNN- and Long-Short-Term-Memory (LSTM)-based AE with an attention mechanism. The learned latent representation was used for supervised classification, optimizing a joint reconstruction and classification loss. Applied on the raw EEG signals of the dataset 2a (classification performed on four MI classes) with a slicing window approach (window length of 400 time points and a step size of 50), the proposed method reached an accuracy of 61%. No information regarding the application of a bandpass filter was given. Notably, it showed improved performance with limited training data, highlighting its suitability for short calibration scenarios. (Sartipi and Cetin, 2024)

Zhang et al. (2021) investigated transfer learning for subject-independent BCIs using the Deep ConvNet of Schirrmeister et al. (2017). Different fine-tuning schemes were compared, ranging from adapting only the classification layer to retraining the entire model. Using a LOSO setup with subsequent subject-specific fine-tuning, the best performance was achieved by adapting only the final layers, while full retraining led to overfitting. The results indicate that early network layers capture subject-independent features and that transfer learning can improve performance with limited data while significantly reducing training time. (Zhang et al., 2021)

#### 2.1.4 Microstate Approach

Liu et al. (2017) were the first proposing a microstate approach for classifying MI tasks. They conducted a microstate analysis for left-hand and right-hand MI EEG data (taken from the dataset 2a of the BCI Competition IV) by comparing multiple parameters of the obtained microstate sequences of both MI tasks. Discriminant features were indicated by a paired t-test and fed into an SVM. In their microstate analysis, Liu et al. (2017) obtained the four canonical microstate templates from the clustering algorithm. The microstate sequences of the EEG trials were built using the software Cartool (Brunet et al., 2011). By comparing the parameters mean duration, time coverage ratio, occurrences per second as well as the transition probabilities across the two tasks, they found significant differences in the first three mentioned parameters for microstate A and B as well as diverse transition dynamics except for the transitions from microstate C to D and vice versa. Those parameters were thus used as input features for an SVM. Classification was performed on each subject individually (subject-specific BCI approach). Liu et al. (2017) reported that their approach yielded higher accuracies compared to conventional feature extraction methods with a mean classification accuracy of 89.17% and a standard deviation of 8.1.

### 2.2 Classification Approaches on Microstate Sequences

To the best of our knowledge, the temporal structure of microstate sequences has note been exploited for MI classification in the context of BCIs. Since this is one of the main methodological contributions of this study, we describe the approach of Sikka et al. (2020) in greater detail and revisit it in the context of MI classification in section 3.4. Sikka et al. (2020) investigated the temporal dynamics of microstate sequences from resting-state EEG data, recorded before and after stress induction. Four microstate templates were obtained using the modified k-means algorithm in a grand-mean clustering across all subjects. Backfitting was performed at Global Field Power (GFP) peaks only with interpolation inbetween peaks. The obtained microstate sequences were further converted to one-hot matrices. Besides the original representation of the microstate sequences, Sikka et al. (2020) also tested an intermediate representation to condense the encoding scheme. An AE with an encoding RNN-layer consisting of 40 LSTM units and a decoding layer mirrored in structure was used to perform a reconstruction of the microstate sequences. Based on the latent representation, a shallow Dense Neural Network was attached to classify the data to pre and post stress condition. The training of the AE and the classifier was performed jointly, thus backpropagating the reconstruction as well as the classification loss simultaneously. The findings of Sikka et al. (2020) support the assumption, that microstate sequences contain temporal information, as random data led to significantly worse reconstruction results. Overall, the reconstruction accuracy decreased with increasing length of the data to be reconstructed (200-2000 ms). As the intermediate, thus shorter, representation did not lead to higher performances for longer sequences, the limitation cannot be attributed to the capacity of the RNN but rather to a limited memory effect of the temporal dynamics in microstate sequences. In contrary, the classification accuracy increased for longer input sequences from 67% for 200 ms up to 73% for 1600 ms, highlighting the long-range dependencies. Furthermore, it is interesting to point out that the authors could not find any significant differences between the pre and post stress data when analyzing the conventional microstate analysis parameters (mean duration, time coverage, occurrences and transition probabilities), which is in contrast to the findings of the study of Liu et al. (2017).

## 3 METHODS

### 3.1 Data Description

In the present work the dataset 2a from the BCI Competition IV was used. EEG data was recorded from nine subjects performing four different MI tasks, namely left- respective right-hand movement, the imagination of movement of both feet as well as the tongue, indicated by a cue appearing on the screen in front of them. For each subject, two sessions on two different days were recorded, the former being the training dataset and the latter the evaluation dataset for the BCI competition. Each session consists of six runs, each containing 48 trials (12 per MI task), resulting in a total number of 72 trials per task per session. (Brunner et al., 2008)

A trial was set up as follows: each trial started with a fixation cross appearing on the screen along with a short acoustic tone (*t* = 0 s). The cue was presented after two seconds (*t* = 2 s) by an arrow pointing left, right, down or up (referring to the four classes left-hand, right-hand, feet or tongue). After 1.25 s the cue disappeared and the subjects were instructed to carry out the MI task until the end of the trial (*t* = 6 s), where the fixation cross disappeared. The trials were separated by short breaks. The recording was performed with 22 Ag/AgCl electrodes placed across the scalp following the international 10-20 montage standard, whereby the left mastoid was used as reference and the right mastoid as ground. The EEG signals were bandpass-filters between 0.5 and 100 Hz with a sampling frequency of 250 Hz. Furthermore, three electrooculography (EOG) channels were included for artifact removal. For this purpose, each session started with a measurement of the EOG influence comprised of a recording with open eyes, closed eyes and eyes movement, taking approximately five minutes in total. In this work, only the classes ’left-hand movement’ and ’right-hand movement’ were used.

### 3.2 Microstate Analysis

#### 3.2.1 Data Preprocessing

The raw EEG data was first bandpass filtered in a 1-40 Hz band and average re-referenced. To remove the physiological eye-movement artifacts, Independent Component Analysis (ICA) was utilized using the EOG channels as reference channels, after which a final bandpass filter has been applied. Two different ranges for bandpass filters were tested: one ranging between 8-15 Hz, adapted from (Liu et al., 2017), and one ranging between 8-30 Hz. The former can be seen as a small upper extension of the mu rhythm (usually defined between 8-13 Hz), the latter comprises the entire alpha and beta rhythm bands. Both frequency ranges are mentioned in literature as relevant ranges for analyzing MI or motor execution data (Pfurtscheller and Da Lopes Silva, 1999; Pfurtscheller et al., 2005; Lapenta and Boggio, 2014). All steps described above were performed in Python using the MNE library.

#### 3.2.2 Microstate Clustering

The microstate clustering was performed by submitting spatial maps at time points of local maxima of GFP to the modified k-means clustering algorithm. Four and five cluster centers were extracted on each subject individually and additionally, a grand-mean clustering on the combined data of all subjects was conducted. For each clustering, the global explained variance (GEV) as

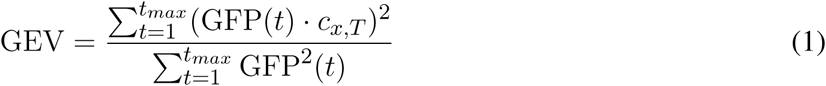

with *c_x,T_* being the spatial correlation between the prevailing topography **x** and the microstate template **T** that best matches the prevailing topography at each time point *t*, was calculated. (Murray et al., 2008)

When clustering with four microstates, all subjects show two microstate templates that highly match with the two canonical microstates A and B. The two remaining templates could in some subjects be corresponded to the canonical forms C and D, however high inter-subject variability was observed. The grand-mean clustering leads to the canonical forms A-C as well as an additional template with a clear left-right orientation, named microstate E in the further work, as depicted in Figure 1. When setting the number of microstates to five, the canonical forms A-D were present in seven subjects along with microstate E. Although reduced when working with five microstate templates, the inconsistency over subjects could still be observed with the used data from the BCI competition IV and therefore corresponds with the findings from Kleinert et al. (2024). As a conclusion, this work uses the grand-mean templates for five microstates extracted from the data of all subjects for the further procedure with a GEV of 77%. The grand-mean clustering was performed on the data of session 1 and successfully reproduced for session 2, resulting in the microstate templates shown in Figure 1. For the microstate clustering, the Python library Pycrostates was used (Férat et al., 2022).

**Figure 1.**
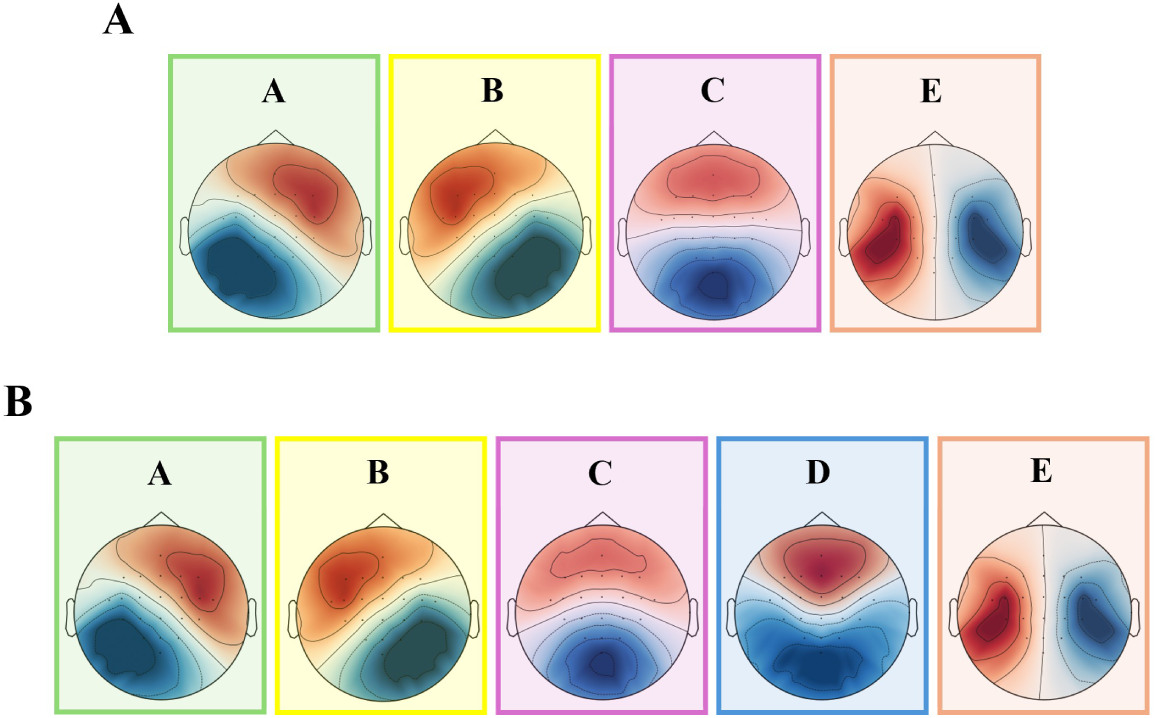
Obtained microstate templates for the grand-mean clustering over the entire EEG data from all subjects. **(A)** Microstate A, B and C can be recognized from the canonical forms. Microstate D can not be found, but instead a microstate E occurs, with a clear left-right orientation. **(B)** All the canonical forms A-D are present as well as microstate E.

#### 3.2.3 Microstate Segmentation

While the clustering had been performed on the continuous EEG data, the segmentation was executed on the extracted epochs. The time span of an epoch was set to 4 s with *t*_min_ = 2 s and *t*_max_ = 6 s around the stimulus (left- or right-hand movement). The segmentation was performed following the continuous approach, thus backfitting at each time point. The open-source Python library Pycrostate was used, which implemented the smoothing algorithm by enabling the user to define the degree of smoothing, the half-window-size as well as the minimum segment length, according to Pascual-Marqui et al. (1995). A grid search was performed to find the optimal parameters of the smoothing procedure taking the average global map dissimilarity (GMD) at GFP peaks as a score function. GMD that quantifies the similarity between the observed topography **x** and a template **T** was thereby defined as

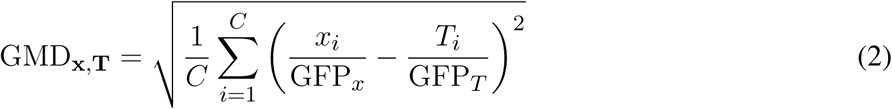

(Murray et al., 2008), where normalization by GFP for each electrode value in a channel, with *C* denoting the number of channels, ensures that differences in global field strength were disregarded. Polarity invariance was achieved by considering both positive and negative orientations and selecting the minimum dissimilarity. The minimum segment length was set to 40 ms and the smoothing factor and half-window-size were ranging from 1-3 respective 2-6. The best parameter combination had been found for a factor that equals 1 and a half-window-size of 5, which was then applied for the segmentation of epochs of left- and right-hand movement. An overview of the complete segmentation process can be seen in Figure 2.

**Figure 2.**
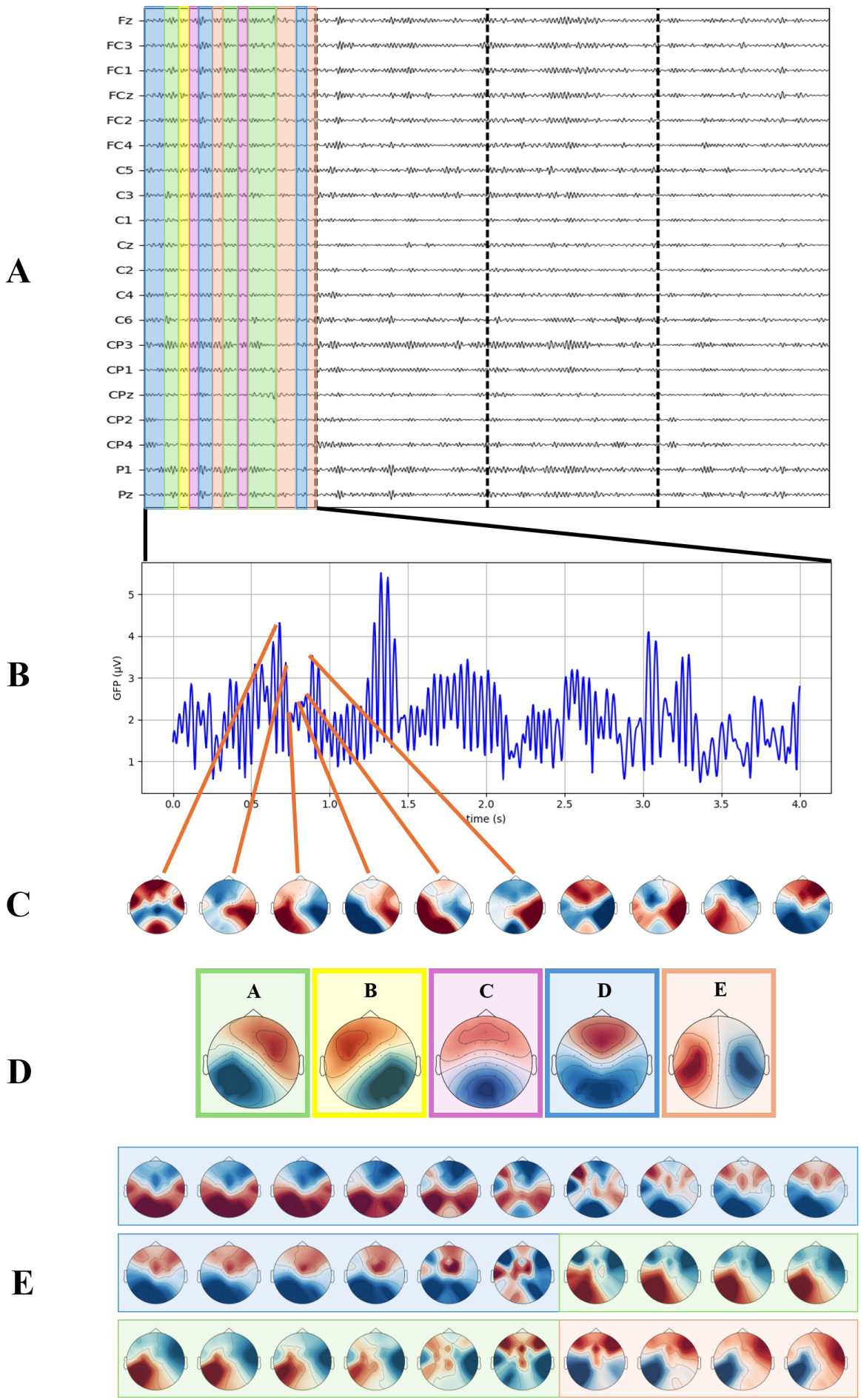
Microstate segmentation method (adapted from (Michel and Koenig, 2018)). **(A)** Continuous EEG signal. **(B)** GFP curve of the first 4 seconds of the shown EEG signal. **(C)** Topographies at times of consecutive GFP peaks that are fed into the clustering algorithm. **(D)** Resulting templates from the modified k-means clustering, labeled A-E. **(E)** Backfitting or segmentation: assignment of the templates to the continuous EEG signal based on the highest correlation and considered smoothing factors. Microstate segments are color-coded in **(A)**.

#### 3.2.4 Microstate Parameter Analysis

Based on the obtained microstate sequences from all left- and right-hand epochs, the characteristic static parameters mean duration, occurrence rate, and time coverage, describing the average duration, frequency, and relative presence of each microstate, respectively, were computed using the Pycrostates library. In addition, transition probabilities represented in a transition matrix, were analyzed to capture the temporal dynamics (microstate syntax). The parameters received for the two conditions were compared and statistically tested for significance to identify possible triggers, thus input features, for classification. The microstate analysis was performed on the data from session 1 only, to not involve inter-session variability. The results of Liu et al. (2017) could not be replicated, as none of the examined parameters reveal significant differences between left- and right-hand MI trials that were consistent across subjects, which is why a classification approach using DL methods was conducted.

### 3.3 Baseline for Classification

In the present work, a classification using conventional methods was conducted to validate reported results on the dataset 2a before comparing them with the proposed approach using microstate sequences. Towards this end, CSP was utilized to extract features from the preprocessed EEG signals and bandpass filtered to 8-30 Hz. The extracted features were used as input into an SVM. Epochs were extracted from *t*_min_ = 2.5 s to *t*_max_ = 4.5 s, as in the winning algorithm of the BCI Competition IV for this dataset (Ang et al., 2008). The CSP class from MNE was implemented to extract six pairs of spatial filters that were further logarithmic transformed before feeding them into the SVM with linear kernel function. Both the subject-specific and the subject-independent approach were performed using the data of session 1. The subject-specific models were once tested on the hold-out data from session 1 and once on the data from session 2. For the subject-independent models, LOSO was applied for performance measuring.

### 3.4 Classification Approach based on Microstate Sequences using Deep Learning Methods

#### 3.4.1 Preparation of Data

Microstate sequences, available as a string of labels from A to E, were first converted to one-hot matrices and then sliced into shorter time series. Thereby, different sequence lengths were tested varying from 200 to 4000 ms (representing a complete epoch from *t*_min_ = 2 s to *t*_max_ = 6 s). The slicing was done in a sliding-window manner with a step size of 10 time steps (40 ms) to increase the number of labeled data samples. All sliced segments from one complete trial kept the same target label, as they have originated from the same MI task, either the imagination of the left or the right hand moving.

#### 3.4.2 Reconstruction of Microstate Sequences

Following the study of Sikka et al. (2020), in a first step an LSTM-based AE was trained to reconstruct the microstate sequences. The model architecture was adopted from the study and consists of a single encoding RNN-layer and a single decoding RNN-layer, each built out of 40 LSTM cells. The encoder of this unsupervised learning architecture leads to a latent representation, which is then used by the decoder to reconstruct the input sequence. As the latent representation holds relevant structure of the input data but reduced in dimension, it is suitable for the subsequent classification task. The model architecture was regularized by introducing a dropout rate of 0.2 into the LSTM layers. While Adam with a learning rate of 0.001 as optimizer had been adopted from (Sikka et al., 2020), Categorical Cross-Entropy was chosen as loss function, in contrary to (Sikka et al., 2020), where Mean Squared Error was utilized as loss. The AE was trained on microstate sequences obtained with a bandpass filter of 8-15 Hz and 8-30 Hz for the sequence lengths 200, 400, 1000, 2000, 3000 and 4000 ms. A train-test split was performed to evaluate the reconstruction performance. The models were each trained on 500 epochs to ensure convergence. Early stopping (patience parameter fixed to 30) and a validation split of 0.2 were set.

#### 3.4.3 Feature Extraction and Classification

After the stand-alone reconstruction of the microstate sequences had been examined on their performances over different sequence lengths, a classifier had to be introduced into the model architecture to perform the desired discrimination between left- and right-hand MI trials. A 2-step approach, where the AE is first trained to come up with a learned representation, which is in a second step fed into a classifier, as well as an end-to-end approach by training reconstruction and classification loss simultaneously were tested in the present study.

##### 3.4.3.1 2-Step Approach

For the 2-step approach, the latent representation from the LSTM-based AE was fed into a small Dense Neural Network, composed of two fully connected layers with 64 respective 32 units. The final binary classification was performed by a sigmoid classification layer. Dropout was utilized as regularization and Binary Cross-Entropy served as loss function. To compare the influence of the varying sizes of the model input on the classification results, the classifier was trained for each of the above defined sequence lengths over 100 epochs using a validation split of 0.2. The models were trained each with a train-test split performed on all trials from all subjects, once for session 1 and once for session 2.

##### 3.4.3.2 End-to-End Approach

The end-to-end approach was carried out using a sequence length of 2000 ms. For the classifier architecture, batch normalization layers were introduced in between the fully connected layers, resulting in a total number of 20525 trainable parameters for the AE and a further 4929 for the classifier head. The model was trained by backpropagating the combined loss, consisting of the reconstruction as well as the classification loss, as visualized in Figure 3. Categorical Cross-Entropy served as reconstruction loss and was weighted with a factor 0.2. For the classification loss Binary Cross-Entropy was used and *β* = 1 applied, as the classification performance should be preferred during training. The pre-trained AE from the stand-alone reconstruction task for a sequence length of *S* = 500 served for initializing the combined model by adopting the initial weights. For the first 10 epochs of the training, the weights of the encoder part were freezed to further emphasize the classification tasks. The combined models were trained over 100 epochs, using Adam as optimizer with a learning rate of 0.001.

**Figure 3.**
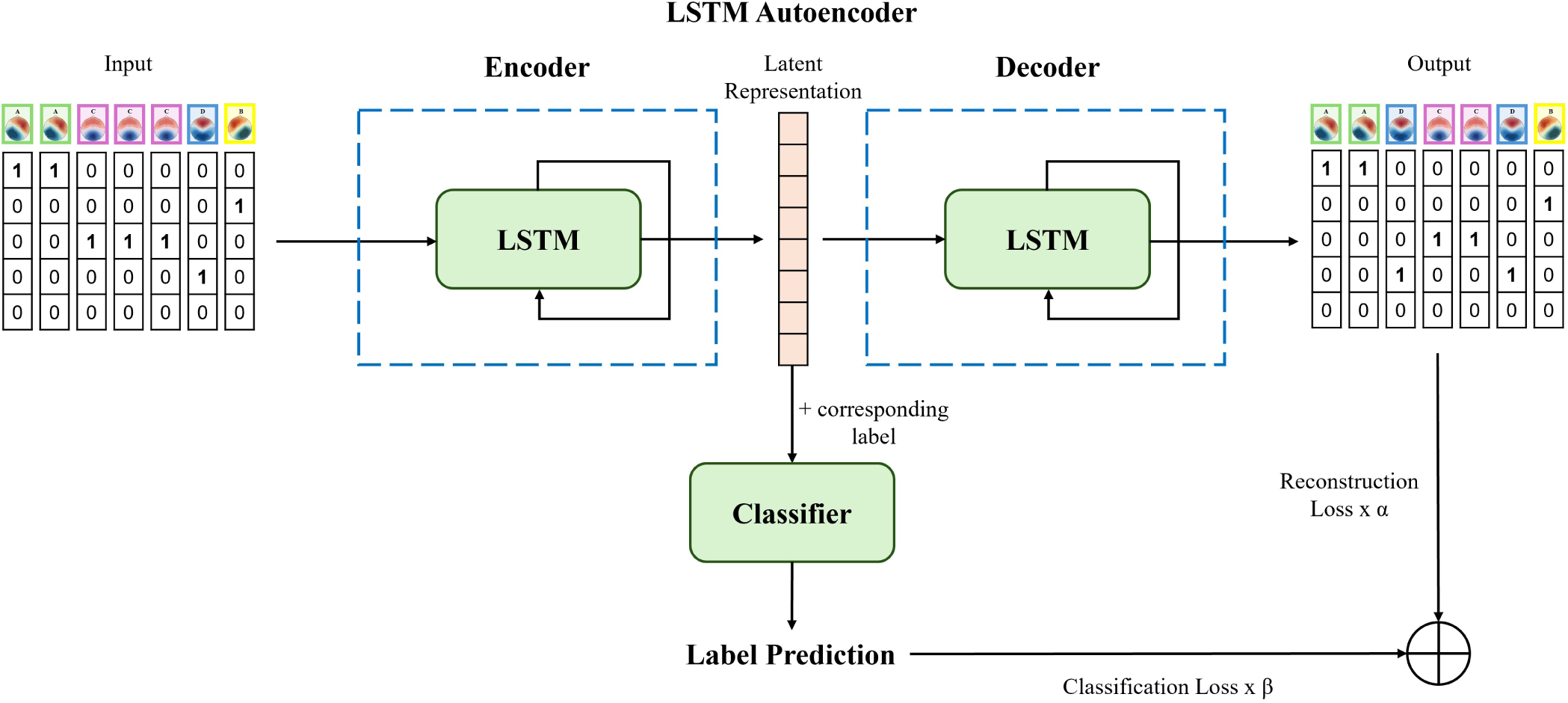
Model architecture of end-to-end training: reconstruction and classification loss are back- propagated jointly. *α* was set to 0.2 and *β* to 1 to emphasize the classification task over the reconstruction.

In total, the end-to-end approach using the semi-supervised architecture was performed in three different experimental setups. In a first setup, the data of all subjects was used jointly to indicate the general performance of the combined model. In a second and third setup, a subject-specific as well as a subject-independent classification were conducted to compare the classification results with existing state of the art approaches. Each of the three setups was examined once with microstate sequences obtained when bandpass-filtering the EEG data to 8-15 Hz as well as when using a broader filter from 8-30 Hz. To first exclude the effect of session-to-session variance, only the data from one recording session was used for training and testing the models. The other session was held out for cross-validating the models’ performances on session-to-session transfer.

##### 3.4.3.3 End-to-End Manner Using All Subjects’ Data

In a first step, the MI EEG signals from all subjects were jointly used for training the combined model. Thereby, a balanced train-test split ensured that all subjects were distributed equally in the training and testing data, using 70% of the trials of each subject for training and the remaining 30% for testing. For further analysis reasons on the session-to-session transfer, the described train-test split was performed on session 1 of the BCI Competition IV and the obtained model was cross-tested on session 2 and vice versa.

##### 3.4.3.4 Subject-Specific Classification

For the subject-specific approach, the train-test split was performed on each subject individually. Therefore, the resulting training size for each subject-specific model is reduced in comparison to when using all subjects’ microstate sequences. The train-test split was first performed on the data of session 1. To further test the session-to-session transfer, the trained models from session 1 were tested on the unseen data from session 2.

##### 3.4.3.5 Subject-Independent Classification

LOSO served as the method for subject-independent classification. Thereby, the obtained microstate sequences from all except for one subject were used for training a model and the remaining subject was used for testing.

#### 3.4.4 Transfer Learning

Two different transfer learning approaches have been examined in the present study, namely the re-training of a subject-specific model on the same subject’s data from a second session (facing session-to-session transfer) as well as the fine-tuning of a subject-independently trained model on data from an unseen subject. Transfer learning was performed by re-training or fine-tuning the classifier part of the model architecture whereas the weights of the AE were not further adopted. Towards this end, a stand-alone classifier with the same architecture as in the combined model was built and initialized with the pre-trained weights from the subject-specific respective subject-independent model. The classifier was fed with the latent representations that were obtained when applying the microstate sequences to the pre-trained encoder part of the combined model. This approach was preferred to re-training the original entire model with the LSTM-layers frozen, due to the reduced computation time^1^.

For both approaches, different learning rates were tested, as proposed by Zhang et al. (2021). Besides the learning rate of 0.001, which was used for training the subject-specific and subject-independent models, the decreased learning rates of 0.0005 and 0.0001 were included in the study. Bearing in mind the convenience for users in a real-world scenario, which requires short calibration times and thus only a small number of necessary EEG recordings, the classification performance for a new subject was analyzed for different sizes of data samples that were used for re-training.

## 4 RESULTS

### 4.1 Reconstruction of Microstate Sequences

The reconstruction accuracy was assessed for each trained AE model for the sequence lengths *S ∈ {*50, 100, 250, 500, 750, 1000*}*. The blue curves in Figure 4 show that the reconstruction performance decreased rapidly with increasing sequence length. While the reconstruction reached about 98% for microstate sequences of 200 ms, the accuracy dropped to 25% when using sequence lengths of 4000 ms, thus an entirely extracted MI epoch (from *t*_min_ = 2 s to *t*_max_ = 6 s).

**Figure 4.**
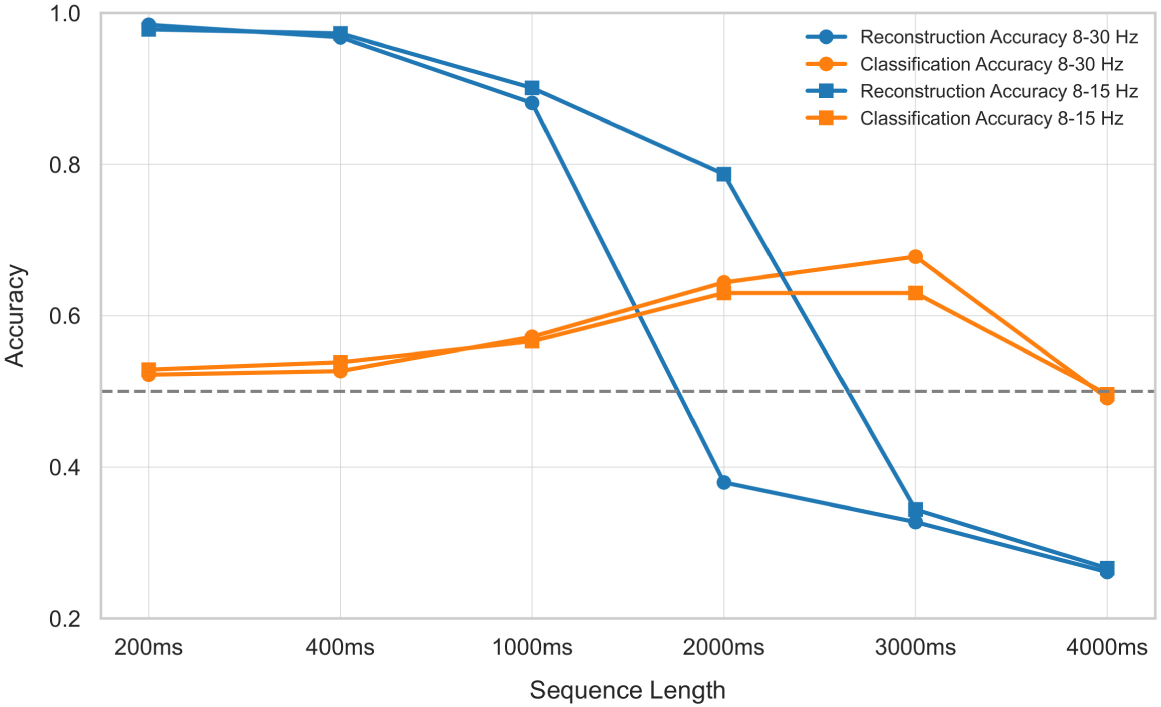
Reconstruction and classification accuracy over different sequence lengths when using a 2-step approach for the bandpass filters of 8-15 respective 8-30 Hz: first, an AE is trained for reconstructing microstate sequences. The latent representations of the AE are fed into a classifier in a second step. Chance level of classification marked at 0.5 with dashed line. Note that the chance level of reconstruction with a 5-dimensional input-vector lies at 0.2.

### 4.2 Feature Extraction and Classification

In the following, the results of the classification task are stated. First, the 2-step approach is reported, followed by the end-to-end approach, where the results are clustered in the subject-specific and the subject-independent classification. Finally, the results of the transfer learning investigations are presented.

#### 4.2.1 2-Step Approach

Figure 4 illustrates the reconstruction (blue curves) and classification accuracy (orange curves) over the range of examined sequence lengths when training the semi-supervised model in a 2-step approach for the two bandpass filters. The reconstruction shows optimal performance for short sequence lengths, whereas classification accuracy increases with longer sequence lengths. A maximum in classification performance is reached at microstate sequences of 2000 respective 3000 ms with about 63% respective 68%. When using the entirely extracted MI epoch of 4000 ms, classification accuracy drops back to chance probability, as visible in Figure 4. It can be seen that a sequence length of 2000 ms, thus *S* = 500, shows a good compromise between reconstruction and classification performance.

#### 4.2.2 End-to-End Approach

Training the semi-supervised model for sequence lengths of 2000 ms in an end-to-end manner on the data of all subjects led to a classification accuracy around 78% for both session 1 and 2. When testing the trained model on the data of the respective hold-out session, classification accuracy drops back to chance level. For both the 2-step approach as well as the end-to-end approach, a t-SNE was utilized to visualize the learned representation of the AE; however, no interpretable patterns regarding the MI task label could be identified (results not shown).

#### 4.2.3 Subject-Specific Classification

The subject-specific microstate approach was performed using batch normalization layers and in an alternative setup without. A considerable performance gain or drop with batch normalization could not be observed. Nevertheless, the presented results refer to the models applying normalization, as the weights from these models were taken for initializing the classifier neural networks for the transfer learning approach.

Table 1 shows the results of the different subject-specific classification approaches on dataset 2a of the BCI Competition IV. The displayed accuracies of the proposed microstate method refer to the setup of bandpass filtering to 8-30 Hz, as this filter led to better performances than the narrower filter of 8-15 Hz. The microstate approach is compared to the baseline results using CSP and a SVM as well as the state of the art accuracy in (Lotte et al., 2009), where a FBCSP was utilized. The results are grouped by the data they were tested on: once testing within session 1 and once testing on data from session 2. The last column shows the results of the session-to-session transfer learning approach. It can be seen that classification based on microstate sequences with the semi-supervised neural network performs better in comparison to the baseline that uses CSP for feature extraction and a SVM as classifier. This performance gain of the proposed method is observable when testing the models on data originating from the same session as the training data as well as when testing on MI trials recorded on a different day. Furthermore, it can be seen that the microstate approach performs well on a subject-specific level when all trials originate from the same session, with a mean accuracy of 92%, but cannot perform well in the session-to-session transfer as the mean accuracy drops back to chance level (*M* = 48%). The same effect was found for the baseline model. The state of the art performance with a mean classification accuracy of 81%, as reached in (Lotte et al., 2009), cannot be achieved without further re-training on the new session’s data. These findings are consistent with the results when training on all subjects’ data, where a session-to-session transfer was also not possible. The transfer learning results in Table 1 are obtained when training on 70% of the subject’s data from the second session and tested on the unseen 30% with a learning rate of 0.001. The achieved accuracy of 89% exceeds the state of the art value and indicates transfer learning as a promising approach.

**Table 1.**
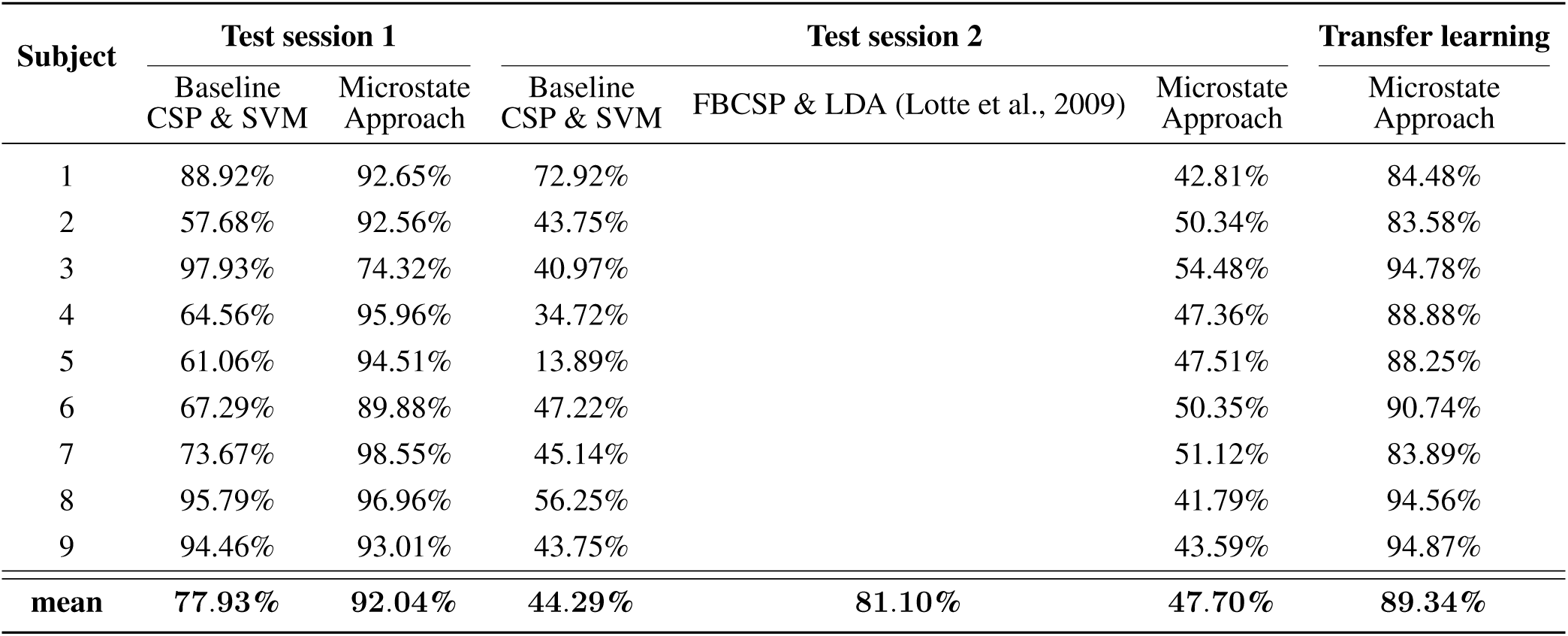
Accuracies of subject-specific approaches: comparison of baseline model with CSP as feature extraction method, a state of the art FBCSP approach, and the proposed microstate approach. The results are grouped by the data they were tested on, with the difference that on the one hand, the test data originated from the same session that the model was trained on and on the other hand, from a second session. The last column shows the results for the transfer learning approach on the data of the second session. Note that for the FBCSP approach in (Lotte et al., 2009), only the mean accuracy is available.

| Subject | Test session 1 |  | Test session 2 |  |  | Transfer learning |
| --- | --- | --- | --- | --- | --- | --- |
|  | Baseline CSP & SVM | Microstate Approach | Baseline CSP & SVM | FBCSP & LDA (Lotte et al., 2009) | Microstate Approach | Microstate Approach |
| 1 | 88.92% | 92.65% | 72.92% |  | 42.81% | 84.48% |
| 2 | 57.68% | 92.56% | 43.75% |  | 50.34% | 83.58% |
| 3 | 97.93% | 74.32% | 40.97% |  | 54.48% | 94.78% |
| 4 | 64.56% | 95.96% | 34.72% |  | 47.36% | 88.88% |
| 5 | 61.06% | 94.51% | 13.89% |  | 47.51% | 88.25% |
| 6 | 67.29% | 89.88% | 47.22% |  | 50.35% | 90.74% |
| 7 | 73.67% | 98.55% | 45.14% |  | 51.12% | 83.89% |
| 8 | 95.79% | 96.96% | 56.25% |  | 41.79% | 94.56% |
| 9 | 94.46% | 93.01% | 43.75% |  | 43.59% | 94.87% |
| mean | 77.93% | 92.04% | 44.29% | 81.10% | 47.70% | 89.34% |

#### 4.2.4 Subject-Independent Classification

Table 2 compares the results for the subject-independent classification approaches on dataset 2a of the BCI Competition IV. The values for each subject refer to the LOSO method, where the models are tested on one subject and beforehand trained on the other eight subjects. The accuracies listed for the proposed microstate approach are obtained with a bandpass filter of 8-30 Hz and when using batch normalization layers in the classifier head of the combined model.

**Table 2.** Accuracies of subject-independent approaches: comparison of baseline model with CSP as feature extraction method, the state of the art approaches using FBCSP with a multi-resolution decomposition (Lotte et al., 2009), a shallow-mirror transformer model (Luo et al., 2023) and a semi-supervised deep architecture (Sartipi and Cetin, 2024) (see also section 2.1.3), and the proposed microstate approach. The last column shows the transfer learning results when re-training the subject-independent models on the data of the left-out subject. Note that in (Lotte et al., 2009), only the mean accuracy is available and in (Luo et al., 2023), the classification was performed on the four MI tasks of the dataset.

| Subject | Baseline<br>CSP & SVM | FBCSP & LDA (Lotte et al., 2009) | Shallow-mirror<br>transformer (Luo et al., 2023) | Semi-supervised<br>Deep Architecture (Sartipi and Cetin, 2024) | Microstate<br>Approach | Transfer Learning<br>Microstate<br>Approach |
| --- | --- | --- | --- | --- | --- | --- |
| 1 | 62.50% |  | 78.47% | 58% | 55.09% | 85.34% |
| 2 | 54.86% |  | 56.94% | 68% | 67.05% | 87.25% |
| 3 | 85.42% |  | 88.54% | 68% | 72.06% | 90.38% |
| 4 | 60.42% |  | 61.46% | 47% | 66.37% | 89.43% |
| 5 | 47.92% |  | 50.69% | 71% | 62.88% | 88.75% |
| 6 | 52.08% |  | 66.32% | 62% | 53.94% | 91.79% |
| 7 | 52.78% |  | 53.82% | 51% | 50.38% | 87.57% |
| 8 | 93.06% |  | 81.60% | 68% | 55.46% | 93.24% |
| 9 | 84.03% |  | 67.71% | 53% | 60.47% | 90.93% |
| mean | 65.90% | 70.99% | 67.28% | 61% | 60.41% | 89.41% |

Results are compared to the baseline results using CSP along with three state of the art approaches, namely the FBCSP approach using a multi-resolution decomposition (Lotte et al., 2009), the shallow-mirror transformer model in (Luo et al., 2023) together with the semi-supervised deep architecture consisting of a CNN- and LSTM-based AE with an attention mechanism and a classifier head (Sartipi and Cetin, 2024) (see section 2.1.3 for a more detailed description of the listed state of the art approaches). Note that the latter conducts the classification on all four MI classes of the dataset, whereas the other approaches only use the left- and right-hand MI trials.

Results show that the baseline values using the conventional CSP feature extraction method are within the range of the state of the art accuracies with a mean accuracy of about 66%. The multi-resolution decomposition FBCSP approach in (Lotte et al., 2009) yielded the highest mean accuracy of 71%. The proposed microstate approach reached a mean accuracy of 60%, indicating that the subject-independent models contain some generalized information regarding the class separation. Nevertheless, the challenge of the across-subject transfer is evident. When comparing the microstate approach for the subject-independent classification in Table 2 with the microstate approach for the subject-specific classification with testing on the data of session 2 (see Table 1), the subject-independent approach yields a higher performance, although both setups are tested on data from an unseen subject respective unseen recording session. This might be due to the amount of available training data, as the subject-independent models are trained on eight times the amount of data.

As in the subject-specific classification, the transfer learning results in Table 2 are achieved by using 70% of the hold-out subject’s data for fine-tuning the models and testing on the remaining 30% with a learning rate of 0.001. A mean accuracy of 89% was reached, which signifies a considerable increase in performance compared to the state of the art methods.

### 4.3 Transfer Learning

Further examination on the learning rate of the transfer learning approaches showed that a decreased learning rate also led to a performance decrease. Table 3 shows the mean classification accuracies for the across-session transfer, conducted on both bandpass filters when varying the learning rate. Filtering to 8-30 Hz and keeping the learning rate at 0.001 led to the best accuracy of around 89%. The mean results for the across-subject transfer with a bandpass filter from 8-30 Hz are reported in Table 4. It is visible that the mean classification accuracy from all nine folds decreases from 89% for a learning rate of 0.001 to 77% when decreasing the learning rate to 0.0001. The analysis of the impact of the number of available data samples on model performance revealed that model accuracy increases as the number of training samples grows. Figure 5 illustrates this positive correlation for each subject along with the mean accuracy, which reaches an accuracy of 80% when 2500 training samples are used. This number of training samples can be produced out of 50 EEG MI trials (25 for each task) with the slicing window approach that was utilized in this study and described in section 3.4.1. In other words, the needed calibration time - on average - to initially reach 80% classification accuracy for an unseen subject is around 400 s.

**Figure 5.**
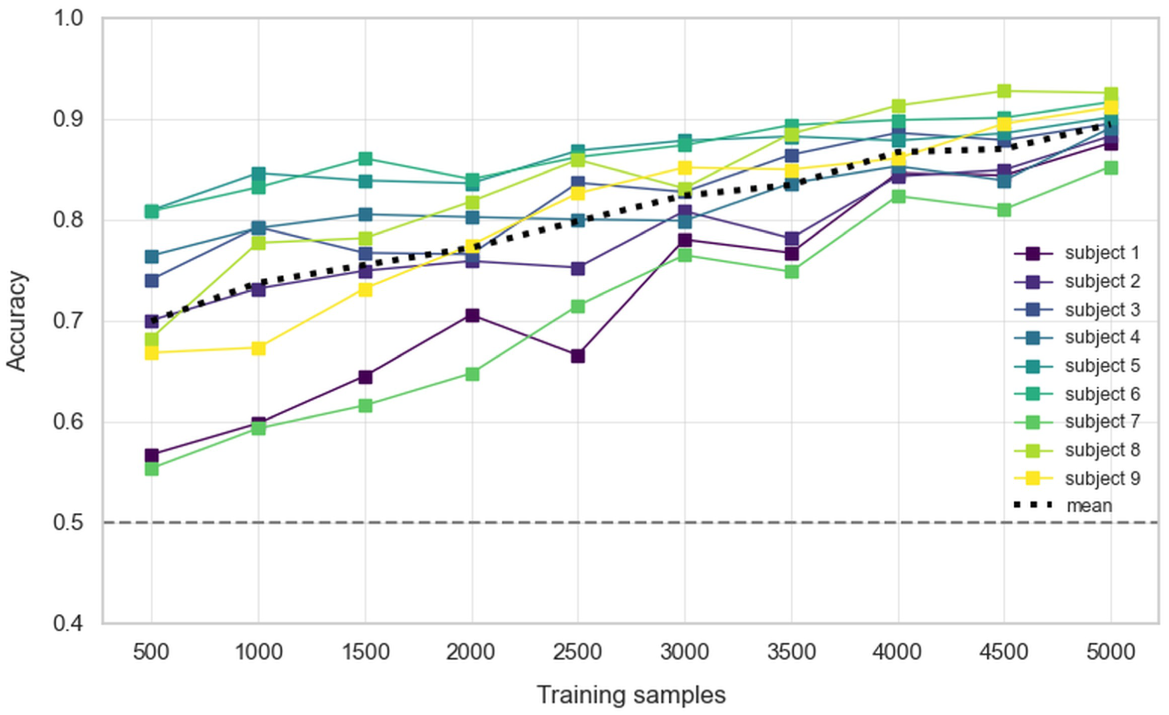
Evaluation of the impact of the number of available training samples on model performance when re-training a model for a new subject. The classification accuracy for each subject and the mean accuracy are displayed. Chance level of classification marked at 0.5 with dashed line. Increasing the number of training samples leads to higher accuracy. On average, an accuracy of 80% is reached when 2500 training samples are available.

**Table 3.** Mean accuracies of transfer learning approach for across-session transfer: comparison on different learning rates when re-training the subject-specific models on the data of the same subject from a second session. Mean accuracies from all nine subjects displayed.

| Bandpass<br>filter | Learning rate |  |  |
| --- | --- | --- | --- |
|  | 0.001 | 0.0005 | 0.0001 |
| 8-15 Hz | 73.22% | 71.79% | 68.23% |
| 8-30 Hz | 89.53% | 88.02% | 75.14% |

**Table 4.** Mean accuracies of transfer learning approach for across-subject transfer: comparison on different learning rates when re-training the subject-independent models on the data of a new subject. EEG data was bandpass filtered to 8-30 Hz. Mean accuracies from all nine folds displayed.

| Learning rate |  |  |
| --- | --- | --- |
| 0.001 | 0.0005 | 0.0001 |
| 89.41% | 87.24% | 77.05% |

## 5 DISCUSSION

In this study, we demonstrated that by employing a semi-supervised DL architecture, consisting of an LSTM-based AE and a Dense Neural Network as classifier, EEG microstate trajectories can be used for classifying left- and right-hand MI when restricting the approach to one recording session and one subject. An across-subject or -session transfer was only possible by re-training the neural network models on the new data, which is in accordance with recent literature on the used dataset. The extent of re-training necessary for reaching a certain classification accuracy was investigated by a transfer learning approach.

A microstate parameter analysis was first carried out with the objective of finding discriminant features as it was presented in the study of Liu et al. (2017). It is of interest that the clustering algorithm on the used dataset with four clusters did not lead to the canonical microstates A-D as introduced by Koenig et al. (1999). This is not in line with the study of Liu et al. (2017), where the typical microstate templates were reported. Reasons might be found in different preprocessing steps, especially regarding artifact handling. In this work, the clustering was performed with five cluster centers, leading to the canonical forms A-D and a fifth microstate E, which exhibits a clear left-right orientation. With the described microstate templates A-E, none of the examined parameters in the microstate analysis revealed significant differences between left- and right-hand MI trials that were consistent across subjects, which is why the present study does not support the previous findings from Liu et al. (2017). The visual inspection of the segmentation or backfitting results revealed the strong influence of the smoothing factors on the obtained microstate sequences. When backfitting according to the highest correlation without smoothing, the microstate segments are in most cases too short to be considered a real microstate. Introducing smoothing parameters into the segmentation process leads to longer microstate segments but at the expense of an increased GMD. As a result, it has to be kept in mind that different smoothing parameters together with the chosen bandpass filter may lead to different microstate sequences and therefore also to different microstate parameters. In this context, the findings of Lehmann et al. (1987) were highly supported. The prevailing topographies particularly matched with one of the microstate templates at time points of GFP maxima and exhibited a high GMD between peaks, indicating transition phases between microstates. Inspired by this result, an alternative approach using only the microstate labels at GFP maxima was tested in this study. As no increase in performance could be yielded, the approach was not further pursued.

It was shown that deep neural networks are suitable for extracting the relevant information and especially RNNs seem appropriate for working with time-courses of EEG microstates. The findings of this study indicate that by employing the proposed semi-supervised model architecture in an end-to-end manner, a latent representation of the EEG microstate trajectories can be found and further utilized for classification. As a compromise between reconstruction and classification accuracy, a sequence length of 2000 ms was chosen. It was observed that the reconstruction accuracy decreased with increasing sequence length, whereas the classification accuracy decreased when using short microstate sequences, which is in line with the findings of Sikka et al. (2020). The latter might be explained by the fact that short segments may be sampled from any part of the EEG trial and therefore may not include the time window containing the task-specific discriminative information.

When applying the proposed approach to a subject-specific classification, taking the training and testing data from one EEG recording session, an average accuracy of over 90% was reached and exceeded the baseline results, where CSP was utilized to extract features from the EEG signals. A session-to-session transfer was not possible without further re-training the model and could therefore not compete with the FBCSP state of the art method in (Lotte et al., 2009). However, when applying transfer learning to the microstate approach to challenge the session-to-session transfer, classification accuracy reached around 90% (see Table 1).

The analysis of the subject-independent classification confirmed previous findings of high inter-subject variance (Sartipi and Cetin, 2024; Lotte et al., 2009; Zhang et al., 2021). As shown in Table 2, the proposed microstate approach achieved a mean classification accuracy of about 60%, which is below the compared state of the art methods (with the highest listed accuracy of about 71% in (Lotte et al., 2009)). Transfer learning applied to the microstate approach, which can be seen as an alternative subject-independent classification approach with short calibration times, led to an average accuracy of about 90%, therefore yielding similar results as on a subject-specific level.

For both the subject-specific and subject-independent models, transfer learning reached around 90% classification accuracy when fine-tuning the classifier heads of the neural networks. Thereby, keeping the learning rate at 0.001 led to the best accuracy, which supports the previous findings of Zhang et al. (2021). An average accuracy of 80% for the across-subject transfer was reached with 2500 training samples for each new subject. This would require 25 EEG trials of each the left- and right-hand MI for a new user. When calculating with eight seconds per trial (orientated at the timing scheme of the dataset 2a from the BCI Competition IV, the necessary data acquisition could be performed in under 10 minutes. Moreover, the time to re-train the classifier with the new data was drastically reduced compared to the training of an entire subject-independent model.

The generally observed high inter- and intra-subject variance supports previous findings on EEG-based BCIs (Aggarwal and Chugh, 2022; Lotte et al., 2018). Although there is a fixed number of microstates that prevail consistent across subjects, the variance between subjects and sessions seems to be reflected in the trajectories and sequences of these microstates. It seems reasonable to assume that each individual brain works differently, so that even the same task, in this case the imagination of the left or right hand moving, evokes different underlying neural activity. More interestingly, the findings of this study indicate a high intra-subject variance, suggesting that, e. g., external factors such as fatigue or attention may also have an important effect on brain activity. Further research could be done regarding the examination of how and to what extent the neural patterns, reflected in microstate trajectories, differ within subjects from day to day. Would it be necessary to adopt and re-train a subject-specific model for each new session? Or would the increasing number of recording days result in the same neural patterns emerging and the model becoming increasingly stable in its classification? These considerations should especially be taken into account regarding a possible real-world application.

Furthermore, differences in the classification accuracy between subjects of around 20% could be observed for the subject-independent microstate approach, that were not further investigated. It could be possible that some of the analyzed nine subjects might not be able to use BCIs, known as ”BCI Illiteracy” (Vidaurre and Blankertz, 2010). An estimate of 15 to 30% of all users are not able to control devices by their brain activity. Vidaurre and Blankertz (2010) named two different groups of BCI illiterates: one, for which a classification algorithm in an offline calibration phase could successfully be trained but no transformation towards an online application with provided feedback, thus control output, was possible, and one, for which no acceptable classification accuracy could be reached during calibration. It is suspected that for the latter, no considerable modulation in the sensorimotor rhythms are evoked during MI (Vidaurre and Blankertz, 2010).

DL approaches, especially transfer learning on pre-trained models, seem highly suitable in the context of BCIs by simultaneously extracting relevant features and using them for classification. This is also reflected in the increasing number of studies regarding DL approaches for EEG-based BCIs, whereof a large proportion use the raw or minimal processed EEG data as input (Roy et al., 2019). As the proposed microstate approach without applying transfer learning did not yield higher subject-independent classification performance than state of the art methods, further research could be done to observe, if transfer learning on a DL model architecture using raw EEG data would reach a similar accuracy. It could be hypothesized that the microstate approach gains an advantage due to the dimensional reduction of EEG microstates compared to raw EEG data. Whereas the raw EEG data with *C* recording channels is available as a *C*-dimensional vector for each time point, the microstate approach reduces the dimension to the number of defined microstate templates.

When integrating EEG microstate features as triggers in a real-world BCI application, additional considerations would be required. A key consideration is the trade-off between classification performance and latency of the live application. While classification accuracy generally improves with increasing microstate sequence lengths, it also entails an increased latency before the application output is available. Furthermore, the frequency with which the classifier evaluates the live recorded EEG data should be assessed. Moreover, the introduction of a threshold between the classification output and the application output requires attention. To avoid uncontrolled or abrupt reactions, the application output should only be triggered when the classification accuracy exceeds a defined threshold, indicating high model confidence.

The investigation of various hyperparameters in the model architecture represents a limitation of the present study. A range of values, e. g., the learning rate, were tested, but mostly confined to the transfer learning approach due to the shorter times needed for the transfer learning model to complete the training. On the one hand, slight adoptions to the proposed model could lead to a further performance gain. On the other hand, the generally observed high inter-subject variance could lead to an individual set of best hyperparameters for each subject, which would be in conflict with the desired ability to generalize.

In the general context of the black box behavior of deep neural networks, visualization may play a decisive role. As proposed in (Sikka et al., 2020), visualization methods may be applied to identify if certain LSTM cells react more pronounced to specific microstate trajectories. In this work, the learned latent representation of the AE was visualized to indicate, if it contains information regarding the classification. No clear segregation or patterns regarding the two classes could be found when visualizing the latent representation by a t-SNE plot. This may be due to the high complexity of the microstate sequences (each composed of 500 5-dimensional one-hot vectors). The lack of effective visualization and therefore the limited interpretability of the neural network highlights another limitation of the present work and points to an important direction for further research.

Despite the aforementioned limitations, the present study signifies the potential of EEG microstate trajectories for extracting BCI triggers. The proposed semi-supervised model architecture showed promising results in capturing information regarding the performed MI task from EEG microstate trajectories. It could further be demonstrated that only a small number of EEG trials is necessary to adopt pre-trained models on data from a new session or subject. High inconsistencies of microstate trajectories between subjects and trials remain a challenge regarding real-world applications and offer scope for further research aiming calibration-free BCIs, e. g., by employing transfer learning approaches.

## GENERATIVE AI STATEMENT

During the preparation of this manuscript, the authors used OpenAI’s ChatGPT-5 for language editing and to improve readability. The authors reviewed and edited all AI-assisted modifications and take full responsibility for the content of the published article.

## AUTHOR CONTRIBUTIONS

AW: Methodology, Formal analysis, Project administration, Software, Visualization, Writing - original draft, Data curation, Validation, Writing - review & editing, Conceptualization, Investigation. MG: Writing - review & editing, Supervision, Conceptualization, Methodology.

## ACKNOWLEDGMENTS

We thank the members of the Brain-Computer Interface and Cognitive Systems Lab, who supported this project. The Cartool EEG Software (cartool.unige.ch) was developed by Denis Brunet at the Functional Brain Mapping Lab, then at the Epilepsy and Networks Lab, University of Geneva, Switzerland, and is supported by the Center for Biomedical Imaging (CIBM), Switzerland.

## DATA AVAILABILITY STATEMENT

The dataset analyzed for this study can be found from the BCI Competition IV.

## Footnotes

1 As the loss in the combined model is composed of the reconstruction as well as the classification loss, the backpropagation has to be performed through all layers, even if the weights of the AE-layers are frozen or the factor *α* for the reconstruction loss is set to 0. This is especially computationally extensive for the LSTM-layers.

